# First characterization of metabolomic and lipidomic exchange over the healthy human brain

**DOI:** 10.64898/2026.06.16.732772

**Authors:** Dario Vrdoljak, Hannah G. Caldwell, Jennifer S. Duffy, Jay M.J.R. Carr, Madden L. Brewster, Alcazar Armando Magaña, Peter Rasmussen, Travis D. Gibbons, David B. MacLeod, Philip N. Ainslie

## Abstract

The energy turnover and metabolic flexibility of the human brain extend beyond a primary reliance on carbohydrates and oxygen. Building on work in anesthetized patients with cerebrovascular pathology, we quantified trans-cerebral arteriovenous differences in healthy humans to isolate and characterize the most abundant cerebral metabolite and lipid species. We observed a net release of acylcarnitine from the cerebral circulation, indicating that these species are active in mitochondrial fatty acid β-oxidation occurring in the healthy resting brain. Furthermore, a strong association was apparent between variability in the brain’s respiratory quotient (RQ) and activity within the purine salvage pathway, particularly with the uptake of guanosine monophosphate. In a larger sample size (n = 210), we further established that biological variability around an RQ of 1.0 - typically interpreted as exclusive carbohydrate oxidation - coincides with coordinated arteriovenous shifts in metabolomic and lipidomic pathways. The variability is not only visible through complex omics pathways, but is also strongly related to the oxygen carbohydrate index of the brain, further supporting that energy substrates other than glucose are exchanged and oxidized across the brain. These findings reveal substantial versatility and redundancy in how the healthy brain maintains its high energetic demands through flexible, interconnected metabolic and lipidomic networks.

## 1. Introduction

In the resting state, the human brain is one of the most metabolically active organ. To maintain basic function, it relies on a stable supply of oxygen and metabolites delivered by active blood flow (Kety & Schmidt, 1945; Scheinberg & Stead, 1949). The most abundant metabolite that the brain uses for energy production is glucose (Attwell & Laughlin, 2001; Fox & Raichle, 1986). Lactate and ketones are well-established alternative fuel sources for the brain that can contribute a meaningful proportion of the brain’s total energy usage when both are made available in circulation (Boumezbeur et al., 2010; Dalsgaard et al., 2004; Hasselbalch et al., 1994; Ide et al., 2000; Mikkelsen et al., 2015; Prins, 2012; Rasmussen et al., 2011; van Hall et al., 2009). However, the brain’s metabolism is not solely related to energy production. For example, many chemical compounds pass from the circulatory system through the cerebral circulation and vice versa, to remove metabolic byproducts, maintain homeostatic balance, facilitate transport and neurotransmission, etc. (Ding et al., 2021; Rasmussen et al., 2021; Wang et al., 2025; Wyss-Coray, 2016). To detect the changes in the pathways of these compounds in humans, one exciting and relatively new scientific approach is metabolomics and lipidomics, which may be applied for hypothesis-driven (targeted or semi-targeted) or hypothesis-generating (untargeted) research (Bird et al., 2026). Metabolomics are a powerful analytic exploration of many (>1000) biochemical metabolites (molecular weight <1,500 Da), which are synchronously examined to gain a unique snapshot of the human metabolome (Clish, 2015). Apart from circulating metabolites, lipidomics can provide additional insight into the brain lipidome by quantifying the changes of individual lipid classes, subclasses and molecular species that reflect metabolic differences (Han, 2016).

Arguably, the gold standard approach for organ-specific metabolomics and lipidomic profiling at the organ level is via the arterial-to-venous differences. For example, the Nobel Prize-winning discovery of the “Cori cycle” included metabolic measures in arterial and venous blood isolated from skeletal muscle and liver (Cori & Cori, 1952). More recently, findings from human brain arteriovenous omics in a cohort of anesthetized patients with different cerebral venous diseases have revealed high net cerebral uptake of glucose, taurine, and hypoxanthine, and significant release of glutamine and pyruvate (Wang et al., 2025). However, the patients were anesthetized and had a variety of vascular diseases, both of which could substantially alter brain metabolism (Slupe & Kirsch, 2018). There are limited studies that have examined arteriovenous metabolomic exchange in the otherwise healthy and resting human brain. In an early and exploratory study, Rasmussen et al. (2010) reported that glucose and lactate are responsible for the majority of carbon exchange across the brain. The authors’ analysis did not reveal a significant export of alternative carbon sources (e.g., carbohydrates or fat) which could explain brain nonoxidative metabolism. An important limitation is the low sensitivity of nuclear magnetic resonance (NMR) spectroscopy for quantifying additional metabolites that may account for the carbohydrate surplus.

The net uptake or release of metabolites and lipids in healthy resting humans may provide new information on the related pathways, as well as provide a critical point of reference and comparison to those with brain injury and/or cerebrovascular pathology (Bird et al., 2026). Therefore, by utilizing sophisticated and validated liquid chromatography (LC) and/or gas chromatography (GC)-mass spectroscopy (MS) approaches (Black et al., 2024; Wang et al., 2025), the primary aim of the present study was to apply trans-cerebral differences to characterize the cellular metabolome and lipidome in healthy humans. The secondary aim was to explore relationships between the arteriovenous differences in the measured metabolites and lipids with well-established metrics of brain metabolism [e.g., oxygen carbohydrate index (OCI), oxygen glucose index (OGI), respiratory quotient (RQ)]. These combined approaches reveal that the normal biological variability of brain RQ around 1.0 – typically interpreted as complete and exclusive carbohydrate oxidation – are explained by coordinated arteriovenous shifts in metabolomic and lipidomic pathways.

## 2. Methods

### 2.1. Ethical Approval

Participants provided informed written consent prior to participating in this study. This study was approved by the University of British Columbia Clinical Research Ethics Board (CREB: H22-01091) and conducted according to the principles established by the Declaration of Helsinki, except for registration in a database.

### 2.2. Omics study (Part 1)

#### 2.1.1. Participants

In total, 12 healthy adults (6 females) completed the invasive arterial-internal jugular venous catheterization protocol. At the time of measurement, their age was 28 ± 5 years, height 175 ± 8 cm, and body mass 69.6 ± 10.8 kg. Participants had no reported history of cardiovascular, cerebrovascular, or respiratory disease and were not taking any prescription medication at the time of their participation. Participants arrived at the laboratory following ≥ 12 hour overnight fast and after having refrained from alcohol and vigorous exercise or activity for ≥12 hours prior to testing. Habitual caffeine users were provided a single black coffee with their standardized breakfast 1 hour prior to the experimental protocol. Data was obtained within the Centre for Heart, Lung and Vascular Health at the University of British Columbia, Kelowna, British Columbia, Canada.

#### 2.1.2. Metabolomics and lipidomics sample processing

To minimize matrix effects and achieve low detection limits, a two-phase extraction was performed as previously described, with modifications optimized for human plasma (Black et al., 2024; Matyash et al., 2008), with modifications optimized for human plasma. Briefly, 50 µL of plasma was homogenized in 300 µL of methanol containing 0.02 mg/mL butylated hydroxytoluene (BHT), followed by the sequential addition of 750 µL of methyl tert-butyl ether (MTBE) and 200 µL of water to induce phase separation. A 500 µL aliquot of the upper organic phase, containing lipids, was collected, dried, and reconstituted in 250 µL of acetonitrile:isopropanol (7:3, v/v) with 0.2 ppm CUDA 12-[(cyclohexylamino)carbonyl]amino]-dodecanoic acid) and 5 µL of SPLASH LIPIDOMIX Mass Spec Standard (Avanti), containing a deuterated mixture of 14 major lipid classes to facilitate identification of elution regions for each lipid class within the chromatogram.

The polar metabolite fraction was collected from a 350 µL aliquot of the aqueous lower phase, dried, and reconstituted in 250 µL of 50% aqueous methanol (v/v) containing 1 ppm internal standards (methionine-d₃, caffeine-¹³C₃, phenylalanine-d_5_, and ferulic acid-d₃), vortexed for 10 min, and centrifuged at 16,000 × *g* for 10 min at 4°C. A 200 µL aliquot of the clarified supernatant was transferred to LC–MS autosampler vials. All samples were stored at −80°C until analysis. A pooled quality control (QC) sample was generated by combining 20 µL aliquots from each sample.

#### 2.1.3. Untargeted Metabolomics LC–MS Analysis

Untargeted metabolomics analyses were performed using a high-resolution time-of-flight mass spectrometer (Impact II, Bruker Daltonics) coupled to a Vanquish UHPLC system (Thermo Fisher Scientific) as previously reported (Black et al., 2024). In brief, chromatographic separation was achieved using an Inertsil Ph-3 column (2 µm, 150 × 2.1 mm) equipped with a matching Ph-3 guard column (GL Sciences). Metabolites were separated using a multi-step gradient consisting of water and methanol containing 0.1% formic acid, with the organic phase increasing from 5% to 99% over 18 min. The column oven was maintained at 55 °C, the flow rate was set to 0.3 mL/min, and the autosampler temperature was held at 4 °C throughout the analysis.

Mass spectrometry data were acquired in both positive and negative electrospray ionization modes using data-dependent acquisition. The capillary voltage was set to 4,500 V in positive mode and −3,500 V in negative mode. The drying gas temperature was 220 °C, and spectra were collected across a mass range of *m/z* 50–1,300. Internal mass calibration using sodium formate was performed prior to each run, resulting in mass accuracy within 2 ppm. Pooled quality control samples were analyzed every twelve runs to monitor instrument performance and analytical reproducibility.

#### 2.1.4. Metabolomics Data Processing and Annotation

Raw LC–MS/MS data were processed using Progenesis QI^TM^ (V3.0.7600.27622, NonLinear Dynamics) with the METLIN^TM^ database integration as previously described with some enhancements (Black et al., 2024). Briefly, to improve data quality, features were filtered using the following criteria: presence of MS/MS spectra, coefficient of variation below 25% in pooled QC samples, signal intensity at least fivefold higher than extraction blanks, and mass accuracy within ±5 ppm. Metabolite identification confidence levels were assigned according to Metabolomics Standards Initiative guidelines (Sumner et al., 2007). Level 1 identifications were confirmed using an *in-house* MS/MS spectral library acquired under matching experimental conditions (Black et al., 2024), while Level 2 annotations were based on spectral matching against public databases including METLIN, HMDB, GNPS, and MassBank. When a metabolite was detected in both ionization modes, the feature showing lower variability in QC samples was selected for downstream analysis. Relative metabolite abundances were calculated from integrated peak areas and normalized across the dataset.

#### 2.1.5. Untargeted Lipidomics LC–MS/MS Analysis

Untargeted lipidomics analysis was performed using a timsTOF Pro 2 mass spectrometer (Bruker Daltonics) coupled to a Bruker Elute LC system. Lipid species were separated on an Acquity CSH C18 column (130 Å, 1.7 µm, 100 × 2.1 mm; Waters) fitted with a CSH C18 VanGuard FIT guard cartridge (1.7 µm, 2.1 × 5 mm), using a gradient adapted from Cajka et al. (2017). Mobile phase A consisted of acetonitrile/water (60:40, v/v) and mobile phase B of isopropanol/water (90:10, v/v), both supplemented with 0.1% formic acid and 10 mM ammonium formate. Elution proceeded as follows: 15% B at 0 min, ramping to 30% B at 2 min, 50% B at 2.5 min, 80% B at 12 min, 99% B at 12.5–13.5 min, returning to 15% B at 13.7 min and held through 17 min. The column was thermostated at 65°C, the flow rate was maintained at 0.5 mL/min, the injection volume was 2 µL, and the autosampler was held at 4°C.

Mass spectrometry data were acquired using trapped ion mobility spectrometry with parallel accumulation–serial fragmentation (TIMS–PASEF) in both positive and negative electrospray ionization modes. In ESI+, the capillary voltage was set to 4,500 V with a dry gas flow of 8.0 L/min; in ESI−, these were adjusted to -3,600 V and 10.0 L/min, respectively. The dry gas temperature was 220°C and the nebulizer pressure 2.2 bar in both modes. The acquisition mass range spanned 150–2,000 *m/z*. Ion mobility separation was performed over a 1/K₀ range of 0.55–1.90 V·s/cm². The PASEF cycle comprised 2 MS/MS scans with a total cycle time of 0.62 s, up to 7 mobilogram peaks per precursor, and active exclusion enabled (release after 12 s) to maximize coverage of lower-abundance lipid species. Ion charge control (ICC) was set to a target of 7.5M counts. External mass calibration was performed at the start of each run by infusing 10 µL of 10 mM sodium formate through a 6-port diverter valve, ensuring mass accuracy within 5 ppm. Pooled QC samples were injected every nine samples to monitor instrument stability throughout the analytical sequence.

#### 2.1.6. Lipidomics Data Processing and Annotation

Raw LC–TIMS–MS/MS data from both polarities were processed in MetaboScape 2024 (Bruker Daltonics) using the T-ReX 4D algorithm, which performs simultaneous alignment across retention time, accurate mass, isotopic pattern, and collision cross section (CCS). Lockmass recalibration with sodium formate constrained mass accuracy to ±5 ppm, and CCS reproducibility was within ±3%. Lipid features were retained if the coefficient of variation (CV) across pooled quality control (QC) injections was <25%, and if mean peak areas in QC samples exceeded those of procedural blanks by at least 10-fold. Lipids were annotated at Metabolomics Standards Initiative confidence Level 2 (Sumner et al., 2007) by spectral matching against MS-DIAL LipidBlast (version 68) libraries (Tsugawa et al., 2015) and using the Lipid Annotation rule-based workflow in MetaboScape 2024 (Lerner et al., 2023).

### 2.2. RQ study (Part 2)

The hypotheses addressed in this study involved retrospective analysis of participants from previously published experiments from 17 studies in healthy humans (Duffy et al., 2026). Eight of these studies (2 unpublished) were data collected from the University of British Columbia, Canada, seven from the Muscle Research Laboratory, Copenhagen, Denmark, and two from the Department of Anesthesiology, Duke University, USA. All studies recruited only healthy adults with no known cardiovascular or neurological diseases between the ages of 19-45 years old (mean 26.4 ± 5.1 years). The current experimental questions involved new objectives and data analysis, which include only resting baseline data, that has not been previously reported. To directly quantify trans-cerebral exchange via the Fick technique, we obtained samples of arterial (brain inflow) and internal jugular bulb (brain venous outflow) blood samples (see below).

All studies were approved by the local ethics committees (University of British Columbia Clinical Research Ethics Board, the Ethics Committee of Copenhagen, and the Institutional Review Board of Duke University Medical Center), carried out in accordance with the Declaration of Helsinki, and informed consent was obtained from all participants. Details on each study can be found in (Duffy et al., 2026).

### 2.3. Testing protocols and equipment

#### 2.2.1. Instrumentation

While resting supine, catheterization of the radial artery and internal jugular vein was performed using sterile technique, under local anesthesia (Lidocaine, 2.0%), and with ultrasound guidance. To accurately isolate cerebral metabolism, it is critical that the blood sampled from the internal jugular vein comprises no contamination from extra-cerebral blood. This was done via cranial advancement (∼15 cm) of the catheter, placing the catheter tip in the jugular bulb. Following this approach, it has been demonstrated to lead to > 97% sampling of venous blood isolated from the brain (Schell & Cole, 2000).

#### 2.2.2. Blood sampling and cerebral metabolism

Arterial and jugular bulb blood samples were drawn simultaneously into pre-heparinized syringes (SafePICO, Radiometer, Copenhagen, Denmark) and analyzed immediately using a commercial blood gas analyzer (ABL FLEX, Radiometer).

For omics analysis, arterial and jugular bulb blood samples were drawn into K2EDTA Vacutainers® (Becton Dickinson, USA). Whole blood was then immediately centrifuged at 600g for 10 min at 4°C. Plasma was aliquoted into cryovials (∼500 µl), flash frozen in liquid N2, and stored at −80°C.

The data acquired from ABL were used in further equations to define brain metabolism. We calculated O_2_ contents from arterial and venous hemoglobin concentrations ([Hb]), saturations (SO_2_) and pressures of oxygen (PO_2_) (Eq. 1).

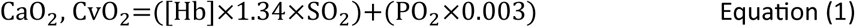

We assessed cerebral CO_2_ parameters by calculating total arterial and cerebral venous CO_2_ content (CCO_2_) per (Douglas et al., 1988). First, we calculated the plasma content of CO_2_ (PCCO_2_) (Eq. 2).

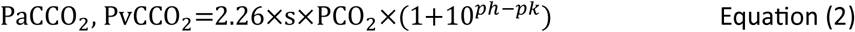

Where *s* is the solubility coefficient of CO_2_ and pK’ is the apparent pK [both calculated per (Kelman, 1966), with the assumption that core temperature was 37°C. PCCO_2_ was calculated for both arterial (PCaCO_2_) and jugular venous (PCvCO_2_) blood using arterial pH, PCO_2_, and pK’ for PCaCO_2_ and jugular venous pH, PCO_2_, and pK’ for PCvCO_2_. CCO_2_ was calculated for both arterial (CaCO_2_) and jugular venous (CvCO_2_) blood, using arterial Hb, SO_2_, and pH for CaCO_2_ and jugular venous Hb, SO_2_, and pH for CvCO_2_ (Eq. 3).

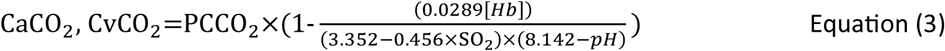

We calculated the respiratory quotient (RQ) of the brain, which was previously determined to approximate 1.0 in resting humans (Lennox, 1931) (Eq. 4). To further examine the metabolic fate of glucose, we employed oxygen glucose (OGI) (Gibbs et al., 1942) (Eq. 5). This value indicates near complete oxidation of glucose: 6 O_2_ + C_6_ H_12_O_6_ (glucose) → 6 H_2_O + 6 CO_2_; approximately 5-10% of the glucose taken up contributes to glycolysis as evidenced by a slight lactate efflux at rest (Dalsgaard et al., 2004; van Hall et al., 2009). Theoretically, OGI values at a 1:1 ratio of oxygen to glucose uptake would equal 6, indicating that all glucose is consumed by oxidative pathways, as the stoichiometry for the oxidation of glucose by oxygen requires one oxygen molecule per each of the 6 carbon atoms in glucose. When OGI decreases below 6, it indicates nonoxidative glucose metabolism is occurring and lactate is dispersed throughout the brain or released into circulation. To account for the lactate efflux the oxygen-carbohydrate index is calculated (OCI) (Eq. 6). If all lactate produced via non-oxidative metabolism is released into circulation, the OCI is 6 (Gibbs et al., 1942).

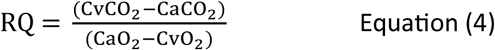

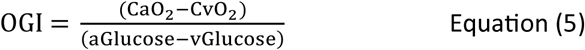

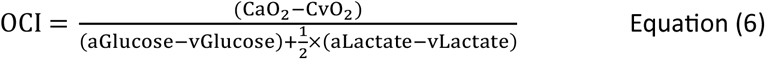

#### 2.2.3. Cerebral blood flow

Volumetric cerebral blood flow (CBF) was measured from the simultaneous diameter and velocity of the right internal carotid artery (ICA) and left vertebral artery (VA) using a 10 MHz multi-frequency linear array duplex ultrasound (Terasmart uSMart 3300, Teratech, Burlington, MA, USA).

### 2.3. Data analysis and visualization

For each participant, we calculated the ratio of metabolite abundance in the vein (V) relative to the artery (A). Using a paired T-test to compare the abundance of these compounds in the artery and the vein, metabolites with *p*-values less than 0.10 were considered to exhibit significant net uptake or release. A mean V/A ratio of less than 1 indicates net uptake, while a ratio greater than 1 indicates net release. To compare data between the artery and vein within each participant, a paired *t*-test was used when the data exhibited a normal distribution (*p* > 0.05 by the Shapiro-Wilk test). Otherwise, the Wilcoxon paired-samples signed rank test was used. To define the difference between magnitudes of net uptake/release, log_2_ was used on V/A ratio values, as well as the magnitude of net uptake/release (Wang et al., 2025). The p-values were tested with the magnitude-based Cohen’s effect size (ES) test with modified qualitative descriptors (trivial ES < 0.2; small ES = 0.21–0.60; moderate ES = 0.61–1.20; large ES > 1.20). Afterwards, p-values were adjusted for multiple comparisons using the Benjamini-Hochberg method, with a FDR cutoff of 0.20. The code and complete analysis pipeline are available in the Supplementary Materials.

Relationships between select variables were analyzed using Pearson’s correlation with the following key variables: RQ OGI, OCI, CBF and GMP. For OGI and OCI, outliers were identified as > upper quartile + 2.2 × interquartile range and < lower quartile – 2.2 × interquartile range (Hoaglin & Iglewicz, 1987); the excluded data corresponded to values considered unphysiological.

All statistical and data analysis were conducted using R version 4.1.2 (Vienna, Austria). The generation of volcano plots, scatter plots, and bar charts was performed in R and GraphPad Prism 10 (GraphPad Software Inc.; San Diego, CA, USA). Metabolite set enrichment analysis and pathway analysis were carried out using MetaboAnalyst 5.0 (Montreal, QC, Canada).

## 3. Results

To characterize the metabolic and lipidomic arteriovenous differences of the healthy resting human brain, we employed a non-targeted analysis of blood samples collected from the radial artery and internal jugular vein. To observe all possible changes from the human metabolome, 7,611 deconvoluted features were measured, from which 170 metabolites were identified using an in-house MS database (Figure 1A, Supplementary Figure 2 and 3) (Black et al., 2024). Similarly, 3,051 lipids features were measured, and 911 were tentatively (L2) annotated according to the MSI guidelines (Figure 3A and Supplementary Figure 1) (Sumner et al., 2007). Annotated compounds were tested for abundance to determine those which are significantly taken up or released by the brain (Figure 1D, 1E, 3D, 3E and Supplementary Table 1). To justify exploration of differences in circulating metabolites and lipids between arterial and venous samples, Partial Least Squares Discriminant Analysis (PLS-DA) and Orthogonal Partial Least Squares Discriminant Analysis (OPLS-DA) were employed (Figure 1B, 1C, 3B and 3C). Both PLS-DA and OPLS-DA demonstrated different metabolic and lipidomic profiles between samples, which led to further analysis of the profiles.

**Figure 1.**
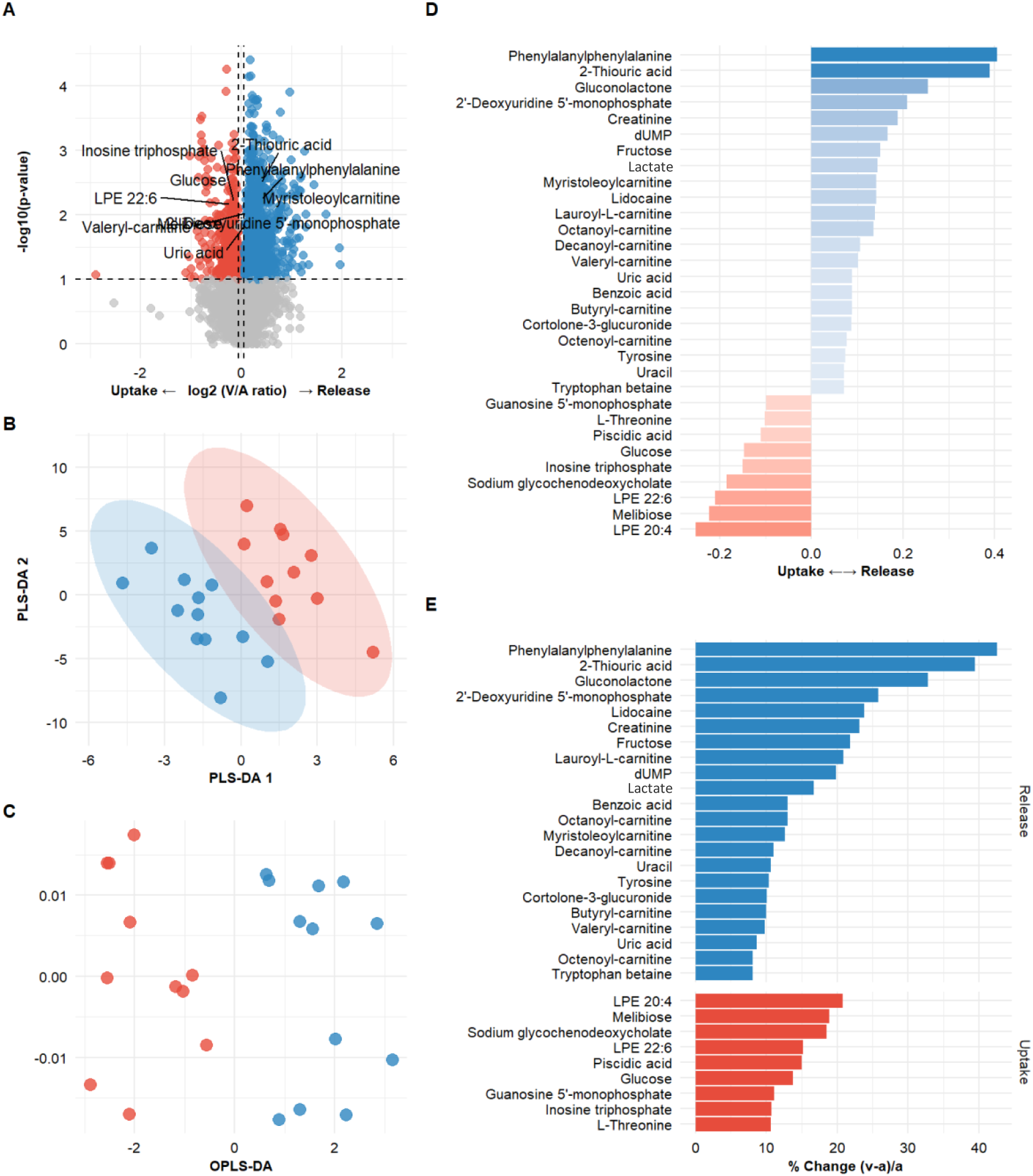
Profiling metabolic exchange between blood and the human brain (A) Volcano plots illustrating metabolites taken up by (left) and released (right). Significance was determined using a two-tailed paired t-test or a one-sample Wilcoxon test. The horizontal dashed line represents p = 0.10 and vertical presents power thresholds set at log2 = ± 0.10. (B) Partial Least Squares Discriminant Analysis (PLS-DA) and (C) Orthogonal Partial Least Squares Discriminant Analysis (OPLS-DA) of untargeted metabolomic data from samples show a difference between arterial (red dots) and venous (blue dots) samples in the same participant. (D) The net uptake and release of metabolites. Bar colors denote metabolite uptake or release, whereas the length of the bars reflects the log2 quantities of net uptake or release. (E) The metabolites taken up and released are represented as a percentage, as calculated via (V-A)/A (Wang et al., 2025).

The data revealed cerebral uptake of lysophosphatidylethanolamine [LPE 22:6 and LPE 20:4 (15-21%)]. Also, high uptake of Glucose (13%) and guanosine-5-monophosphate (GMP) (11%) highlight the metabolic pathways for energy production (Figure 1E). As expected, released metabolites corroborate the uptake compounds. For example, there was a significant efflux of phenylalanylphenylalanine (43 %), 2-Thiouric acid (39 %) and Gluconolactone (33%). Release of these metabolites is consistent with pentose phosphate pathway flux in the brain. Furthermore, the release of lactate (18%) from the human brain is observed in the present data. Acylcarnitines are esters formed between a fatty acyl group and L-carnitine; there was a net release of Carnitines (10-20 %) (Lauroyl-L-carnitine, Octanoylcarnitine, Decanoylcarntine, Valeryl-carnitine, and Butyryl-carnitine). These transport intermediates shuttle activated fatty acyl chains across the inner mitochondrial membrane for β-oxidation (Longo et al., 2016), and are essential for neural metabolism and neurotransmission (Latham et al., 2021) (see Figure 2).

**Figure 2.**
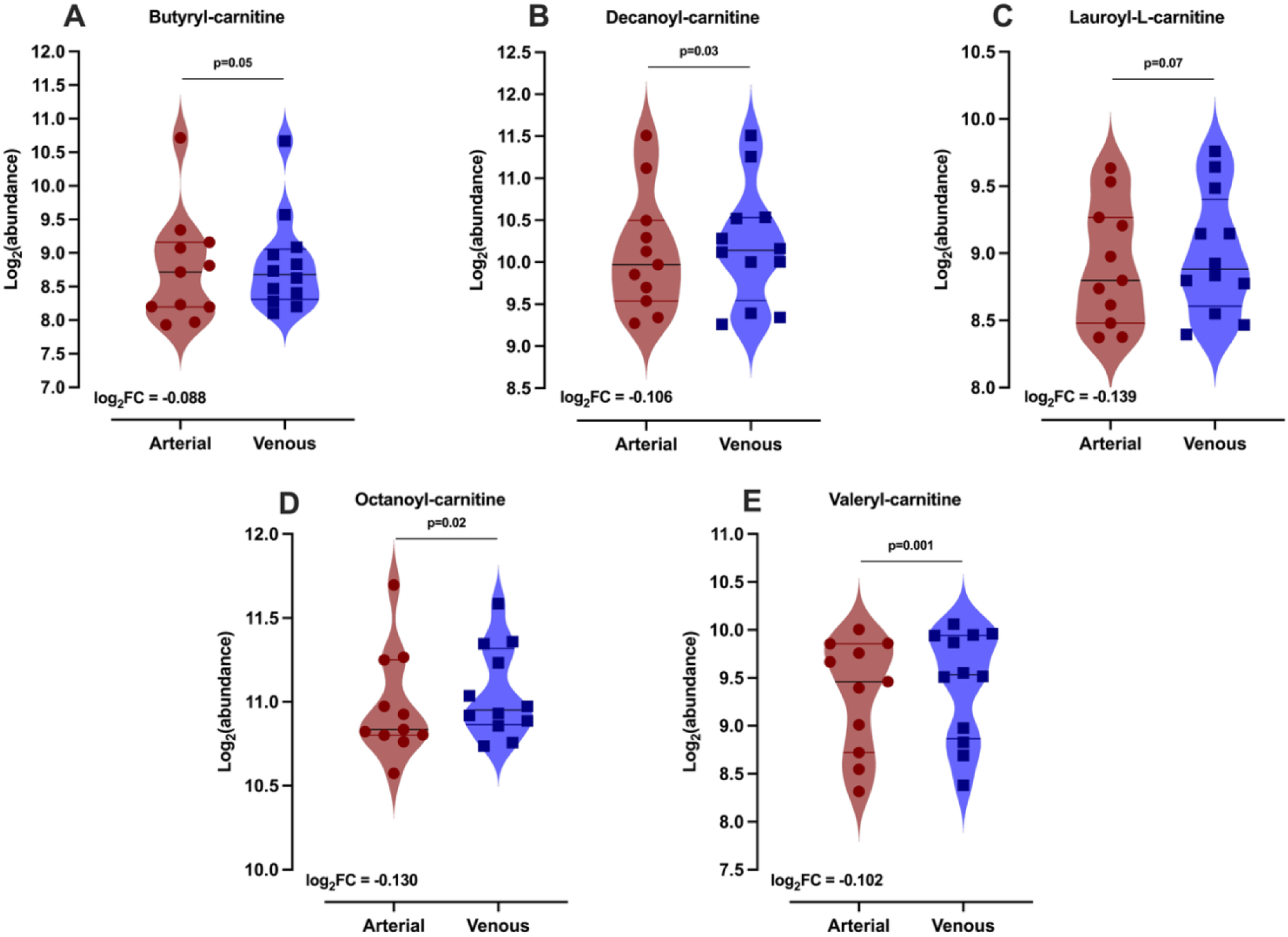
Net cerebral release of five acylcarnitine species by the brains of the 11 subjects: (A) butyrylcarnitine, (B) decanoylcarnitine, (C) lauroylcarnitine, (D) octanoylcarnitine, and (E) valerylcarnitine. The acylcarnitines are depicted separately to further emphasize the novelty behind this finding, as these metabolites serve as obligate transport intermediates for the shuttling of activated fatty acyl chains across the inner mitochondrial membrane for β-oxidation. Violin plots depict the median and lower and upper quartiles of the Log_2_-transformed values. Significance was determined using p-values from a two-tailed paired t-test or one-sample Wilcoxon test.

**Figure 3.**
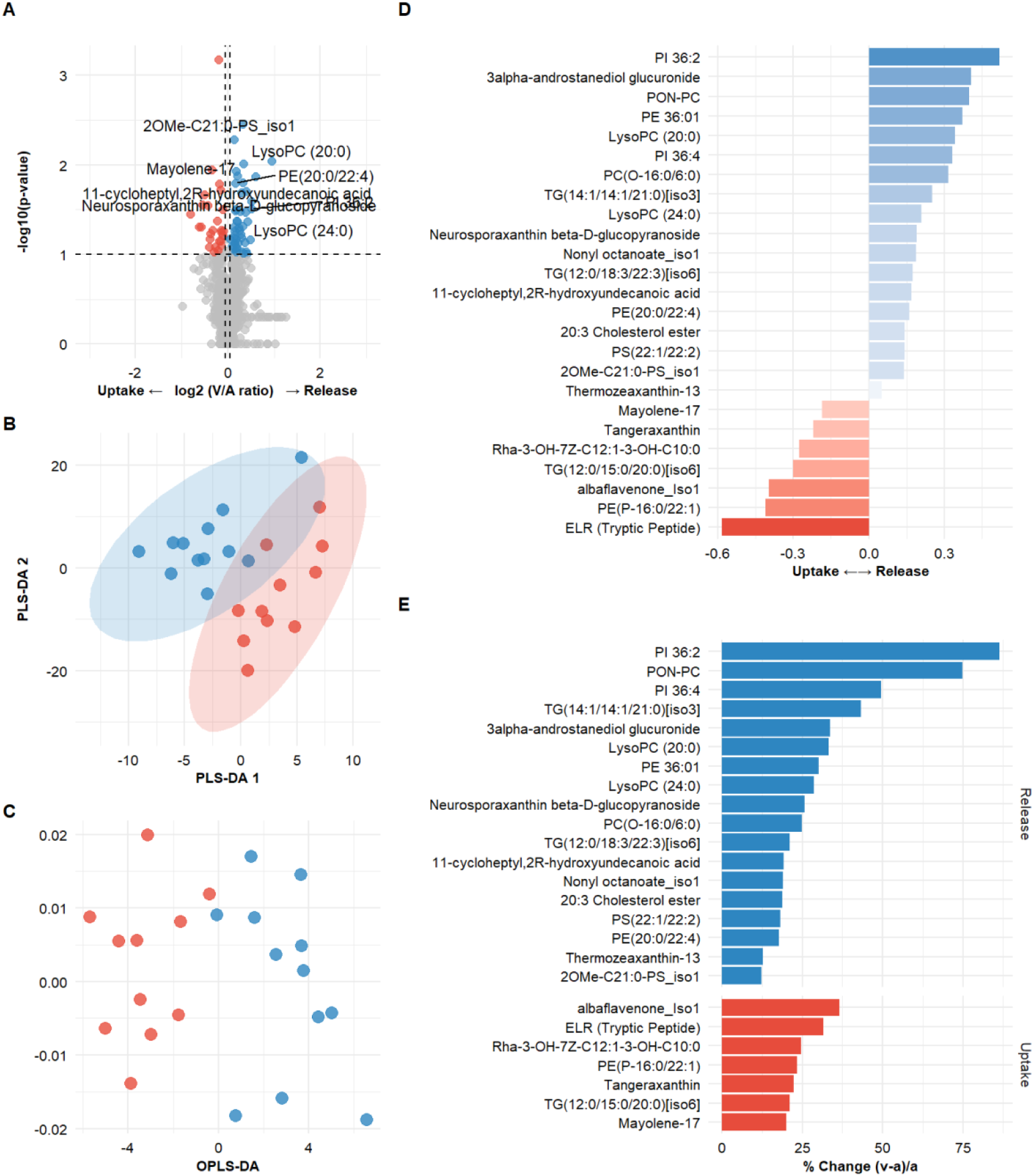
Profiling lipidomic exchange between blood and the human brain (A) Volcano plots illustrating substances taken up by (left) and released (right). Significance was determined using a two-tailed paired t-test or a one-sample Wilcoxon test. The horizontal dashed line represents p = 0.10 and vertical presents power thresholds set at log2 = ± 0.10. (B) Partial Least Squares Discriminant Analysis (PLS-DA) and (C) Orthogonal Partial Least Squares Discriminant Analysis (OPLS-DA) of untargeted lipidomic data from samples show a difference between arterial (red dots) and venous (blue dots) samples in the same participant. (D) The net uptake and release of lipids. Bar colors denote metabolite uptake or release, whereas the length of the bars reflects the log2 quantities of net uptake or release. (E) The lipids taken up and released are represented as a percentage, as calculated via (V-A)/A (Wang et al., 2025).

Apart from circulating metabolites, the cerebral net flux consists of multiple and significantly abundant lipids. Several lipids released from the brain are connected to cell membrane remodelling as well as metabolism maintenance (e.g., Phosphatidylinositol and Phosphatidylcholine [PI 36:2 [83%], PON-PC [75%], and PI 36:4 [36:4]]). In addition, a few different triglycerides were also defined in the lipidomic profile of the brain (TG(14:1/14:1/21:0) [40%] and TG(12:0/18:3/22:3) [23%]), which are mainly connected to neurological function and adaptation. On the other hand, Lysophosphatidylcholines (LysoPC (20:0) [32%] and LysoPC(24:0) [31%]) usually define injury or neuroinflammation. Simultaneously, net influx of lipids consists of peptides (ELR(tryptic peptide) [31%]), and phosphatidylethanolamine (PE(P-16:0/22:1) [21%]), which all serve as neuroprotection, signaling and brain function maintenance.

In the integrated view (Figure 4), we observed that significant pathways corresponding with metabolite circulation are purine, pyrimidine and hexoses metabolism. Lipids were sorted into classes according to the number of corresponding lipids e.g., glycerophospho, sterol, glycero and prenol lipids.

**Figure 4.**
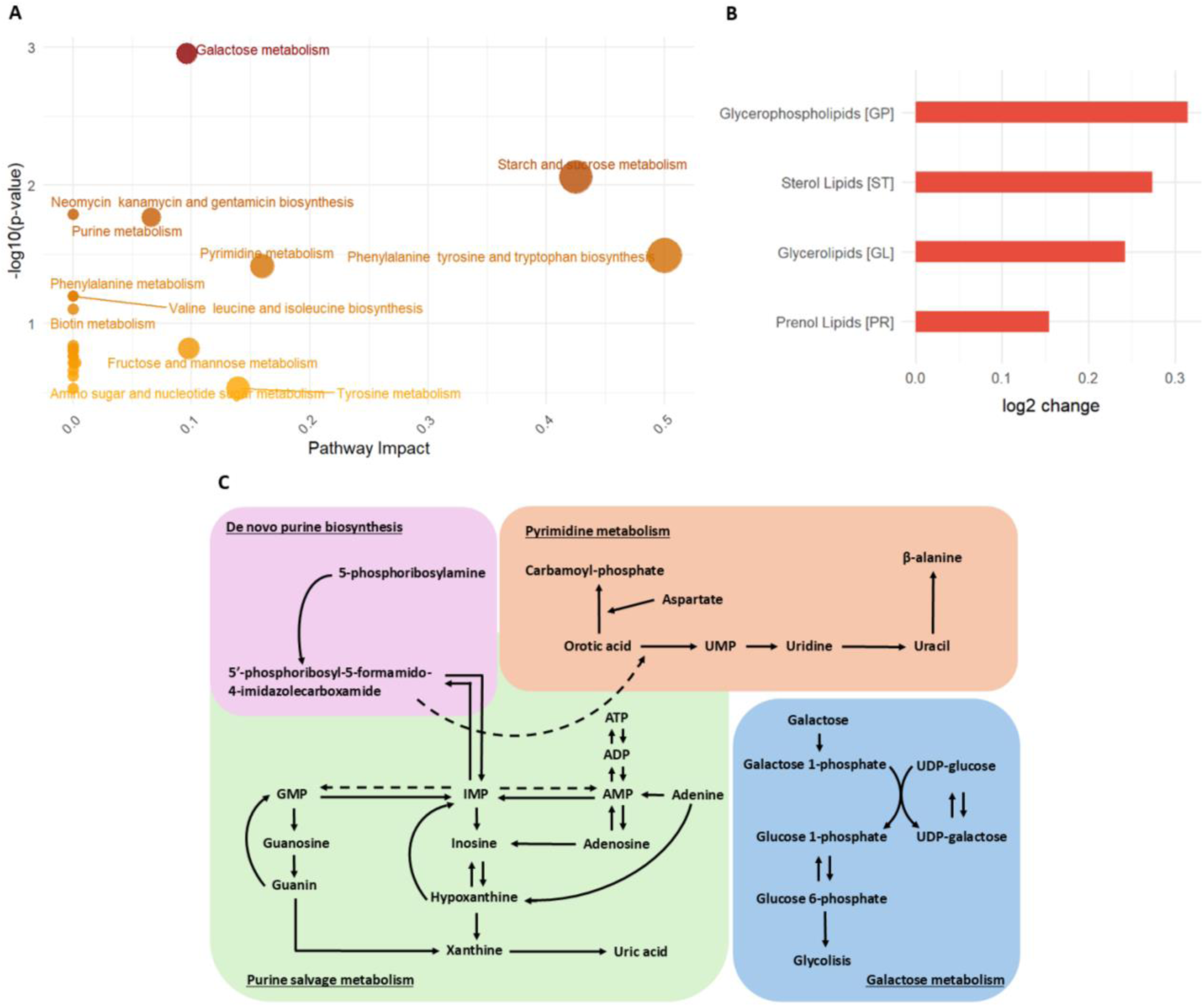
Illustration of significant pathways and classes related to mostly abundant metabolites and lipids (A) Pathway analysis for metabolites exhibiting significant brain A-V differences (p < 0.05). (B) Lipid class as a change in the power of lipids taken up or released by the brain. (C) Summary diagram of representative pathways related to pyrimidine, purine and hexoses (galactose) metabolism in relation to (A).

### 3.1. RQ study

Per Equation 4 and Lennox (1931) the RQ of the brain in normal healthy resting conditions would vary around 1.00. However, as seen from the results of 210 participants, RQ has a range from 0.55 to 1.80. These variations are strongly and positively correlated to OGI and OCI (p<0.001; Figure 5B). A similar and consistent relationship was observed in the smaller data subset from Study 1 (N=11), which was used for further metabolomic analysis (Figure 5A). Further exploration of factors influencing RQ variability showed a significant interaction with Guanosine Monophosphate (GMP) and CBF. As shown in Figure 6, a decrease in RQ relates to higher uptake of GMP (p < 0.001) (Figure 6A) as well as an increase in CBF (p = 0.05) (Figure 5B). Lastly, higher uptake of GMP is related to a higher CBF (p = 0.04) (Figure 6C and Supplementary Figure 4).

**Figure 5.**
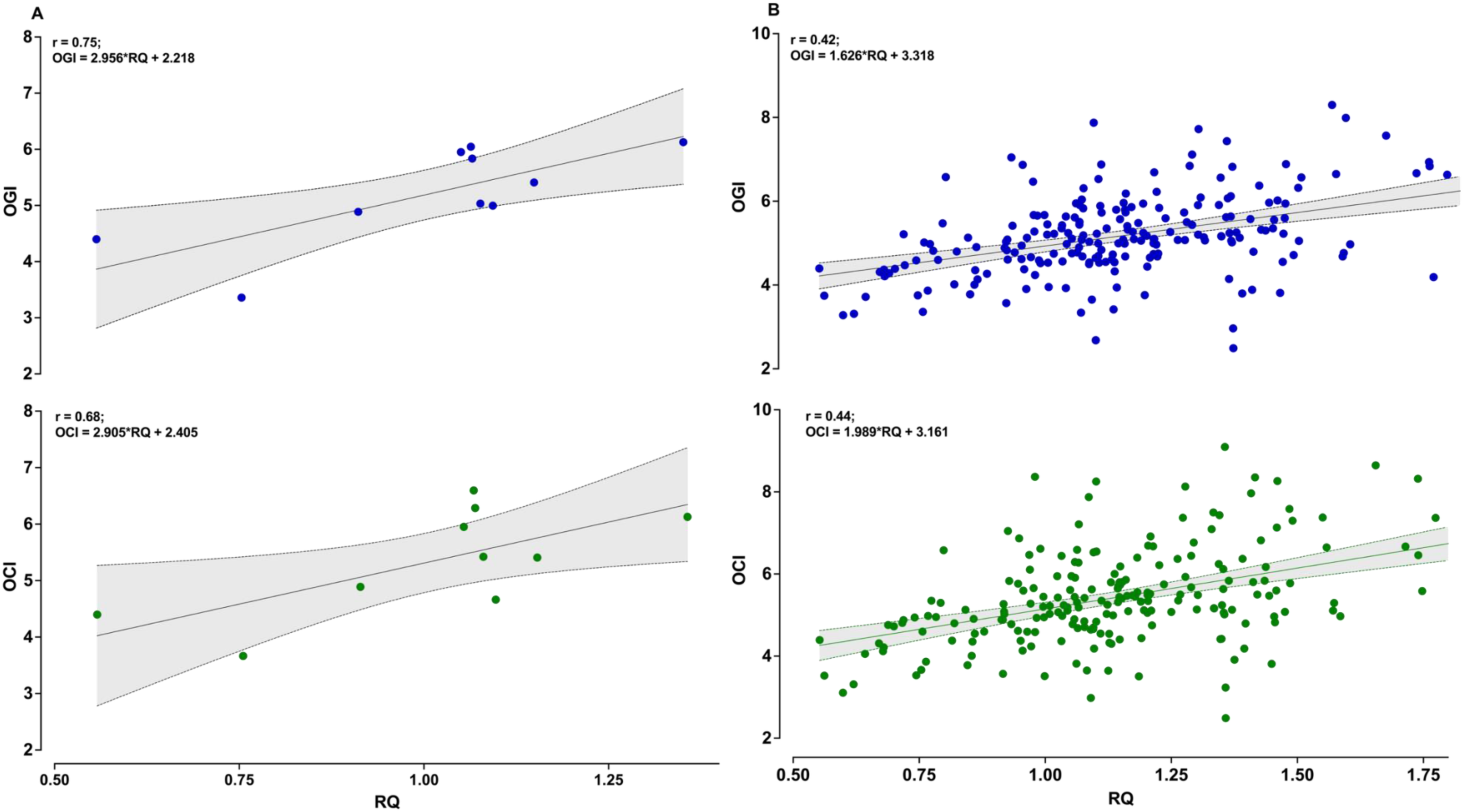
Relationships of oxygen glucose and oxygen carbohydrate index with variations in RQ. (A) Relationship of glucose (OGI) and carbohydrate (OCI) metabolism of the brain with the RQ based on a small sample size (N=11). (B) Depicts consistency of the same relationship between OGI and OCI with RQ in a large sample size (N=210). All results were determined by a linear regression model. Equations of the line and statistics are displayed in each respective panel, while raw data and simple linear regressions depict the trend on each graph. Significance was set at p<0.05; dotted lines represent the 95% confidence interval.

**Figure 6.**
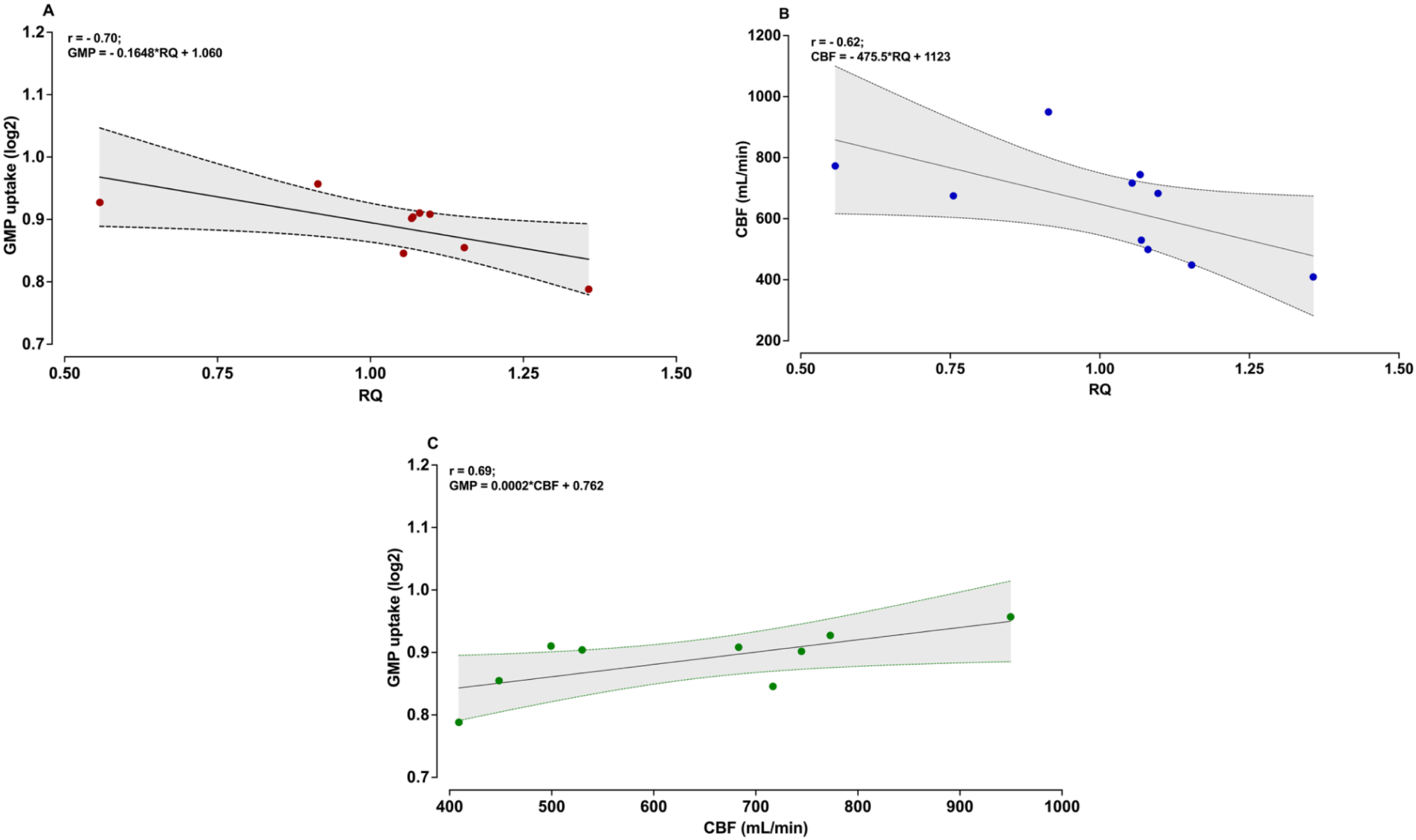
Relationship among CBF, RQ and GMP. GMP magnitude was quantified as log_2_(a-v difference) of signal intensity obtained with LC-MS/MS metabolomic analysis, with positive values representing uptake of certain compound. (A) The magnitude of GMP uptake by the brain is inversely proportional to variability in RQ (p < 0.001). (B) CBF responds to a RQ variability similar to metabolite uptake, showing that when lower RQ relates to higher CBF (p = 0.05). (C) Following GMP and CBF connection to RQ; increase in CBF indicates an increase in uptake of GMP (p = 0.04). The interaction of these factors implicates that variability in RQ is essentially reflected in changes in CBF and related uptake of GMP, leading to Purine salvage metabolism to potentially compensate for a switch towards nonoxidative metabolism and/or lack of substrates in brain. Significance was set at p<0.05; dotted lines represent the 95% confidence interval.

## 4. Discussion

We have quantified trans-cerebral arteriovenous differences in healthy humans to characterize the cerebral metabolome and lipidome. In doing so, we have identified the most abundant and dynamically regulated metabolites and lipid species across the resting human brain. Our findings support a shift from the classical view that the adult brain engages minimally in fatty acid catabolism. This was found in a high net release of acylcarnitine from the cerebral venous circulation, indicating that these species are active in mitochondrial fatty acid β-oxidation occurring in the healthy resting brain. We observed a strong association between variability in the brain’s RQ and activity within the purine salvage pathway, particularly via guanosine monophosphate uptake. Using direct measurements of RQ, we further established that biological variability around an RQ of 1.0 - typically interpreted as exclusive carbohydrate oxidation - coincides with coordinated arteriovenous shifts in metabolomic and lipidomic pathways. The variability is evident not only in complex omics pathways but is also closely linked to the brain’s oxygen-carbohydrate index, also supporting the potential use of alternative substrates as fuel sources. Together, our findings reveal substantial versatility and redundancy in how the healthy brain maintains its high energetic demands through flexible, interconnected metabolic and lipidomic networks. The following discussion highlights the evidence, methodological considerations and the implications of these findings.

### 4.1. Metabolomic and lipidomic characterization of the resting human brain

Arteriovenous brain metabolomic and lipidomic characterization has recently been determined in anesthetized patients (Wang et al., 2025). The results revealed uptake (e.g., lactate and cortisol) and release (e.g., creatine and glutamine) of some metabolites that are consistent with cerebrovascular pathology different than that occurring in the uninjured brain. As acknowledged by the Wang et al. (2025), both pathology (e.g., venous stenosis and/or chronic venous sinus thrombosis) and anesthesia may markedly influence any conclusions on what a ‘normal’ metabolic exchange might be in an otherwise healthy brain. For example, and broadly consistent with the current findings, numerous studies consistently report 5-10% lactate *release* at rest in healthy individuals (Blazey et al., 2018; Dalsgaard et al., 2004; Koep et al., 2025; Rasmussen et al., 2010; Rasmussen et al., 2011). Therefore, by the identification of pathway analysis to define the most significantly expressed metabolites and lipids in the awake and unanesthetized brain, our current findings extend the study of Wang and colleagues (Wang et al., 2025). The main expressed pathways were related to purine and pyrimidine metabolism and were consistently related to one significant metabolite; guanosine 5′-monophosphate (GMP) (Figure 6).

These two (i.e., purine and pyrimidine) metabolic pathways include both the de novo purine biosynthesis and the purine salvage pathway (Figure 4C). However, it is reported that the brain has a limited capacity for de novo purine and pyrimidine synthesis and often relies on nucleosides and nucleotides synthesized de novo in the liver, which are later transferred to the brain to be metabolized through the purine salvage pathway (PSP) (Piero et al., 2011). Despite transfer from the liver to the brain - broadly consistent with the current findings (Fig 3C) - the aforementioned study by Wang et al. (2025), showed PSP expression as one of the most significant pathways. In addition, recent transcriptomic evidence also indicates that the PSP dominates in brain tissue rather than the de novo purine biosynthesis pathway (Frenguelli, 2019; Gessner et al., 2023). Hence, salvage pathways represent relatively rapid and energetically efficient processes that can swiftly increase neuronal activity, particularly during stress situations when substrate need is elevated (Gessner et al., 2023).

The salvage pathway conserves energy by recycling nucleobases derived from dietary sources or nucleotide breakdown, requiring only one ATP per purine synthesized (Murray, 1971). It is reported that PSP relies on two key enzymes acting in parallel: adenine phosphoribosyltransferase and hypoxanthine-guanine phosphoribosyltransferase. Adenine phosphoribosyltransferase converts adenine into adenosine 5′-monophosphate, while hypoxanthine-guanine phosphoribosyltransferase converts hypoxanthine or guanine into inosine 5′-monophosphate or GMP, respectively, inosine can subsequently be converted into adenosine or GMP to yield nucleotide di- or triphosphates (Tran et al., 2024). As illustrated, GMP plays one of the most important roles in this pathway as the last compound before energy production. Notably, the purine salvage pathway in our results was enriched for multiple metabolites that are taken up by the brain (such as inosine and guanosine) and metabolites that are released from the brain (including uracil, uric acid and deoxyuridine monophosphate). Therefore, it seems that the human brain salvages purines (i.e., GMP) from the blood while simultaneously using liver-derived nucleosides to help facilitate its high energetic demands.

Metabolites are not the only compounds circulating through the cerebral vasculature. Specifically, of all human tissues, the brain has one of the highest lipid contents, accounting for 50% of its dry weight (Hornemann, 2021). Brain lipids mainly consist of cholesterol and phospholipids, such as phosphatidylcholine, phosphatidylethanolamine, and sphingolipids (Naudí et al., 2015; Yoon et al.). However, transport of fatty acids across the blood–brain barrier likely involves both diffusional and transport protein-mediated processes. Hence, arteriovenous analysis of these compounds allows for a unique quantification of lipid species and exchange in the brain. Our results showed upregulation in net uptake and release in four main classes of lipids: glycerophospholipids, sterols, glycerolipids, and prenols. These lipids are primarily involved in cell membrane formation, play critical roles in energy storage, the regulation of membrane fluidity and permeability, and signal transduction (Castellanos et al., 2021). Cholesterol, sphingolipids, and glycerophospholipids are abundant in the brain due to the specific flux of lipids regulated by the blood brain barrier (Pifferi et al., 2021). Glycerophospholipids and glycerolipids are the main phospholipid components of cell membranes (Farooqui et al., 2000). Alterations in their composition affect the stability, permeability, and fluidity of neural membranes, which, in turn, lead to neurological diseases (Farooqui et al., 2000). On the other hand, sterols are prevented from crossing the blood brain barrier. For example, there is <1% daily exchange rate between the brain and the periphery; therefore, metabolism of sterols in the brain can be considered almost independent of peripheral tissues (Hartmann et al., 2007; Pifferi et al., 2021). Endogenous cholesterol is mostly generated from astrocytes and, to a lesser extent, transferred to neurons and used for neurite maintenance and synaptic connectivity (Arenas et al., 2017; Naudí et al., 2015). Therefore, normal cholesterol homeostasis is important for maintaining brain function, and its disruption can lead to neurodegenerative and psychiatric diseases, and cognitive deficits, especially in the elderly (Aon et al., 2021; Czubowicz et al., 2019; Lu et al., 2011; Schneider et al., 2017). Understanding the fundamental lipid metabolism in the healthy brain can be used to inform what is normal and abnormal in related derangements with aging and common cerebrovascular pathologies. This information may ultimately be used to guide new biomarker discovery for the development of diagnostic and treatment approaches (Shamim et al., 2018; Vance, 2012).

### 4.2. Acylcarnitine Efflux

Among the most striking metabolomic findings was the net cerebral release of five acylcarnitine species: butyrylcarnitine, valerylcarnitine, octanoylcarnitine, decanoylcarnitine, and lauroylcarnitine, each showing 10 - 20% efflux relative to arterial concentrations (Figure 1D and 2, Table S2). Acylcarnitines are formed by the conjugation of acyl-CoA intermediates with L-carnitine via carnitine acyltransferases and serve as obligate transport intermediates for the shuttling of activated fatty acyl chains across the inner mitochondrial membrane for β-oxidation (Longo et al., 2016). Their net release from the cerebral venous circulation - rather than uptake - indicates that these species are generated within the brain and exported, consistent with active mitochondrial fatty acid β-oxidation occurring in the healthy resting brain.

This observation challenges the classical view that the adult brain relies almost exclusively on glucose oxidation and engages minimally in fatty acid catabolism suggesting that lipid mobilization and fatty acid catabolism are more active in the resting healthy brain than previously appreciated. The translational significance of this finding is considerable. Altered circulating acylcarnitine profiles have been reported in Alzheimer’s disease (Cristofano et al., 2016). Our data now provide a healthy baseline against which such disease-associated deviations can be compared.

### 4.3. PUFA delivery via lysophosphatidylethanolamines

The net cerebral uptake of lysophosphatidylethanolamines containing the two most neurobiologically critical PUFAs - omega-3 docosahexaenoic acid (LPE 22:6, 21%) and omega-6 arachidonic acid (LPE 20:4, 15%) - is particularly noteworthy (Table S2). To date, lysophosphatidylcholines (LPCs) have been the only lysophospholipid class demonstrated to deliver PUFAs across the blood–brain barrier, acting via the sodium-dependent symporter MFSD2A (Alakbarzade et al., 2015; Nguyen et al., 2014). Whether other lysophospholipid classes contribute to cerebral PUFA acquisition has remained unknown. Our arteriovenous data now provide the first *in vivo* human evidence that in resting healthy humans lysophosphatidylethanolamines (LPEs) carrying DHA and arachidonic acid are also taken up by the brain, suggesting that LPEs serve as a parallel PUFA delivery route - potentially through MFSD2A itself, which may have broader substrate specificity than currently recognized, or through complementary transport mechanisms yet to be identified. Given the critical roles of DHA in synaptic membrane composition and neuronal signaling, and of arachidonic acid as the principal precursor for eicosanoid-mediated neuroinflammatory cascades, the preferential uptake of these specific LPE species extends the current LPC-centric model of brain PUFA supply and opens new questions about the molecular determinants of lysophospholipid selectivity at the blood–brain barrier. Notably, PE(P-16:0/22:1), an ethanolamine plasmalogen, also showed 21% net influx, consistent with the known role of plasmalogens as DHA reservoirs and antioxidant membrane components in the CNS (Braverman & Moser, 2012).

### 4.4. Metabolomic and lipidomic insights into changes in oxidative and non-oxidative cerebral metabolism

As seen from our results on 210 resting humans, RQ was related to both OGI and OCI, indicating that the higher the non-oxidative metabolism of carbohydrate (e.g., OGI and OCI < 6), is related to the lower the ratio of CO_2_ release to O_2_ uptake (e.g., RQ < 1). While early work (Gibbs et al., 1942; Kety, 1957) established an RQ of ∼ 1.0 and an expected OGI of ∼ 6 based on complete glucose oxidation, consistent observations of OGI < 6 - indicate that a portion of glucose metabolism is non-oxidative and not fully accounted for by lactate efflux (Blazey et al., 2018; Duffy et al., 2026). Therefore, it remains plausible that stable RQ could be maintained with the uptake of different metabolites, rather than solely glucose and/or lactate. Integrating metabolomic and lipidomic data, our findings suggest that this apparent mismatch is not simply a deviation from glucose oxidation but reflects broader metabolic flexibility in the human brain. Specifically, we found that approx. ∼ 50% of the statistical variability in RQ was related to GMP uptake. Specifically, oxidative metabolism (RQ ∼ 1) and oxidation (OGI ∼ 6) of glucose are reflected in reductions of the GMP uptake (Fig 3), highlighting a potential shift in energy production metabolites. As stated above, the energy-efficient PSP highly relies on the availability of GMP (Gessner et al., 2023). Hence, this relation of RQ and GMP is observed through upregulation of the efficient PSP, which implies periods of deficient availability of energy, where the unique ability to yield additional phosphates in an economical and energetically potent way highlights the importance of this pathway. Hence, we interpret our data to indicate that the variability in RQ may not necessarily reflect pure glucose oxidation, but may also coincide with parallel activity in the purine salvage pathway and/or lipid turnover.

### 4.5. Methodological considerations

A key strength of this study was the gold-standard invasive Fick principle technique to quantify arteriovenous exchange of fuel substrates in vivo in healthy humans (Kety & Schmidt, 1948). The principal assumption of this technique is that jugular venous drainage is solely from the brain. In support of this, it is estimated that 97.3% of jugular venous blood is representative of cerebral sources, such that there is reportedly a small bias (<3%) of contamination with extracerebral blood (Shenkin et al., 1948). Therefore, we believe that IJV sampling was accurate and reliable. Furthermore, despite the existence of these robust relationships between metabolites and variables of cerebral metabolism, it is not possible to determine causation per se with our data. Nevertheless, the consistency of the relationships between OGI and OCI with RQ is apparent across both independent studies. To address this limitation, future work should employ experimental (within-study) manipulation to better test causality (i.e., via the insulin-glucose clamp technique or lipid infusions to target direct manipulations of the metabolite and lipid pathways).

The novelty of the multi-omics approach for the determination of cerebral metabolic pathways in healthy humans must be emphasized; however, we acknowledge that these measures reflect a steady-state “snapshot” in time and do not capture the dynamic flux of the brain. This topic should be addressed in the aforementioned future studies above. The annotation level for many lipid species corresponds to MSI Level 2 confidence. In particular, the assignment of fatty acyl chains to the sn-1 versus sn-2 positions cannot be resolved unambiguously by conventional LC-ESI-MS/MS alone. Therefore, these annotations should be considered putative and interpreted with appropriate caution. Orthogonal structural validation approaches will be important for definitive structural confirmation. For many new or unannotated metabolites or lipids, orthogonal validation is often recommended (Sumner et al., 2007). In the current study, however, the top expressed pathways were consistent with analytical standards (Table Sx). For example, GMP and some carnitine signatures were consistent with certified standards for validated annotations, providing additional reassurances on our data interpretation.

### 4.6. Implications

Our findings highlight the brain’s remarkable metabolic flexibility and the critical role of alternative pathways such as the purine salvage pathway, in maintaining energetic homeostasis. It is uncertain whether glucose hypometabolism is a cause or consequence in the development of neurodegenerative disease (Constantino et al., 2025; Cunnane et al., 2011). However, it is well known that a drop in brain glucose metabolism occurs in advance of cognitive decline (Jagust et al., 2006; Reiman et al., 2004; Small et al., 2000). Therefore, understanding other pathways, such as GMP-driven purine salvage, to provide sufficient energy and maintain “homeostasis” would be important in future human research in both clinical and healthy populations. Beyond the healthy human realm, previous research defined some of the cerebral metabolomic and lipidomic pathways (Wang et al., 2025). In spite of different states of participants (i.e., anesthetized vs. resting; clinical vs healthy), the brain demonstrated similarity in pathway change when it lacked the main substrate (i.e., glucose). In addition to lipids, cholesterol homeostasis is important for maintaining brain function whereby its disruption can lead to neurodegenerative and psychiatric diseases, and cognitive deficits, especially in the elderly. Our results provide new avenues for interpreting metabolomic and lipidomic changes in comparison between healthy and clinical populations, where deviations from these homeostatic mechanisms may underlie disease vulnerability and may ultimately be used to guide new biomarker discovery for the development of diagnostic and treatment approaches.

## 5. Conclusion

The resting human brain is characterized by several important metabolomic pathways and lipidomic classes (e.g., purine and pyrimidine pathways; glycerol and sterol lipids). These pathways generally relate to neuronal protection, membrane remodelling, and signalling. For instance, we see the healthy awake resting human brain actively oxidizes medium-chain fatty acids (evidenced by acylcarnitine efflux) and takes up DHA- and arachidonic-acid-containing lysophosphatidylethanolamines as a parallel PUFA-supply route. Apart from this general characterization, the brain’s versatility in oxidative metabolism is coordinated with shifts in omics pathways that allow such flexibility when sole glucose oxidation fails to provide sufficient energy for the brain’s demands. Together, our findings support a framework in which the healthy brain maintains its high energetic demands through flexible, interconnected metabolic and lipidomic networks.

## 6. Acknowledgments

We are extremely grateful to the numerous (50+) investigators and graduate students who were involved with the extensive data collection for the studies included in this paper. Finally, we gratefully acknowledge Professor Emeritus Niels Secher for his role as investigator and physician in several of the original studies that formed part of this analysis.

